# Transcript-specific induction of stop codon readthrough using CRISPR-dCas13 system

**DOI:** 10.1101/2023.03.08.531701

**Authors:** Lekha E. Manjunath, Anumeha Singh, Debaleena Kar, Karthi Sellamuthu, Sandeep M. Eswarappa

**Affiliations:** Department of Biochemistry, Indian Institute of Science, Bengaluru, Karnataka, 560012, India

**Keywords:** Cas13, CRISPR, nonsense mutation, stop codon, translational readthrough

## Abstract

Stop codon readthrough (SCR) is the process where translation continues beyond a stop codon on an mRNA. Here, we describe a strategy to enhance or induce SCR in a transcript-selective manner using CRISPR-dCas13 system. Using specific guide RNAs, we targeted dCas13 to the downstream region of the canonical stop codons of mammalian *AGO1* and *VEGFA,* which are known to exhibit natural SCR. Results of readthrough assays revealed the enhancement of SCR of these mRNAs (both exogenous and endogenous) caused by dCas13. This effect was associated with ribosomal pausing, which has been reported in several SCR events. Furthermore, our results show that CRISPR-dCas13 can induce SCR across premature termination codons (PTC) in the mRNAs of green fluorescent protein and *TP53*. Finally, we demonstrate the utility of this strategy in the induction of readthrough across the thalassemia-causing PTC in *HBB* mRNA. Thus, CRISPR-dCas13 can be programmed to enhance or induce SCR in a transcript-selective and stop codon-specific manner.

## Introduction

In certain transcripts, ribosomes fail to terminate translation at the stop codon and continue till the next in-frame stop codon, resulting in longer protein isoforms with extended C-termini. This phenomenon called stop codon readthrough (SCR) is mediated by the recoding of stop codons by near-cognate tRNAs or suppressor tRNAs (Schueren & Thoms, 2016). Readthrough isoforms can be different from the canonical isoforms in their function, localization and/or stability (Eswarappa *et al*, 2014; Manjunath *et al*, 2020; Schueren *et al*, 2014; Singh *et al*, 2019). Hence, target-specific manipulation of SCR can have potential applications. For example, SCR of *VEGFA* results in reduced angiogenesis, which can be beneficial in the treatment of cancerous tumours and retinopathies (Eswarappa *et al*., 2014). Furthermore, transcript-specific induction of SCR will have therapeutic benefits in genetic diseases caused by nonsense mutations (e.g., β-thalassemia) that result in premature stop codons in the coding sequences (Keeling *et al*, 2014).

Ataluren, certain macrolide and aminoglycoside antibiotics, and 2,6-diaminopurine are some of the molecules that can induce SCR across premature stop codons (Howard *et al*, 1996; Trzaska *et al*, 2020; Welch *et al*, 2007; Zilberberg *et al*, 2010). Among these, ataluren has received conditional approval from the European Medical Agency for the treatment of Duchenne Muscular Dystrophy. Engineered suppressor tRNAs is another strategy under development for the induction of SCR (Lueck *et al*, 2019). However, all these strategies lack specificity. They can potentially target any stop codon in any transcript, and result in undesired effects. Therefore, there is a need for a strategy that can induce SCR in a transcript-specific manner.

hnRNPA2/B1, an RNA binding protein, binds downstream of the stop codon (9 nucleotides downstream) and positively regulates the SCR of *VEGFA* (Eswarappa *et al*., 2014; Nico *et al*, 2021). Similarly, let-7a microRNA binds downstream of the stop codon of *AGO1* mRNA and enhances its SCR (Singh *et al*., 2019). Even exogenous antisense oligonucleotides that target downstream region of the premature stop codon in *HBB* mRNA can induce SCR (Kar *et al*, 2020). Furthermore, RNA pseudoknot structure formed downstream of the stop codon between the *gag* and *pol* region of murine leukaemia virus induces SCR (Houck-Loomis *et al*, 2011). These observations indicate that transient molecular obstacles downstream of the stop codon can enhance or induce SCR (Manjunath *et al*, 2022). Therefore, targeting of an RNA binding protein specifically downstream of a stop codon can potentially serve this purpose. Here, we have used CRISPR-Cas13 system to achieve this.

RNA targeting Cas13 (formerly C2c2) belongs to the Class 2 Type VI CRISPR-Cas system. This CRISPR-associated enzyme targets RNA in a sequence specific manner with the help of a guide RNA (Watanabe *et al*, 2019). Because of this specificity, multiple applications such as RNA editing, DNA/RNA detection, RNA knockdown and dynamic imaging of RNAs have been developed (Abudayyeh *et al*, 2017; Cox *et al*, 2017; Gootenberg *et al*, 2017; Yang *et al*, 2019). This system has also been used in the diagnosis and the treatment of SARS-CoV-2 infections in a rodent model (Blanchard *et al*, 2021; Freije *et al*, 2019; Patchsung *et al*, 2020). Here, we have applied CRISPR-dCas13 (catalytically inactive Cas13) system to target mRNAs proximally downstream of the stop codon to enhance or induce SCR in a transcript-specific manner.

## Results and Discussion

Since Cas13 can target a specific region of an mRNA with the help of a guide RNA (gRNA), we hypothesised that this system can be used to create a transient molecular obstacle for ribosomes near the stop codon, and thereby, enhance or induce SCR. We tested this hypothesis using the catalytically inactive variant of this enzyme – dCas13, which can bind a specific region of an mRNA without cleaving it (Abudayyeh *et al*., 2017). The gRNAs were designed such that they target mRNAs proximally downstream (4-20 nucleotides) of the stop codon.

### Targeted enhancement of SCR by dCas13

We first tested our hypothesis in *AGO1* mRNA, which encodes Argonaute 1 (Ago1) protein. Ago1 is important for microRNA-mediated repression of gene expression (Meister, 2013). *AGO1* has been shown to undergo SCR across its canonical stop codon resulting in a longer isoform called Ago1x (Ghosh *et al*, 2020; Singh *et al*., 2019). Unlike Ago1 (the canonical isoform), Ago1x cannot repress the expression of target transcripts and serves as an inhibitor of the microRNA pathway when its expression is increased (Singh *et al*., 2019). In cancer cells, Ago1x inhibits dsRNA-induced interferon signalling and promotes cell proliferation (Ghosh *et al*., 2020).

We designed a gRNA such that it guides dCas13 to the region downstream of the canonical stop codon of *AGO1* mRNA (blue region in Fig 1A). A gRNA that does not target any transcript in mammalian cells (non-targeting) was used as control in all assays (Abudayyeh *et al*., 2017). Expression of dCas13 (from *Leptotrichia wadei*) in transfected cells was confirmed by RT-PCR (Fig EV1A). Immunoprecipitation of dCas13 followed by qRT-PCR confirmed the interaction of dCas13 with the *AGO1* mRNA in the presence of the *AGO1*-3′UTR-targeting gRNA (Fig EV1B).

**Figure 1.**
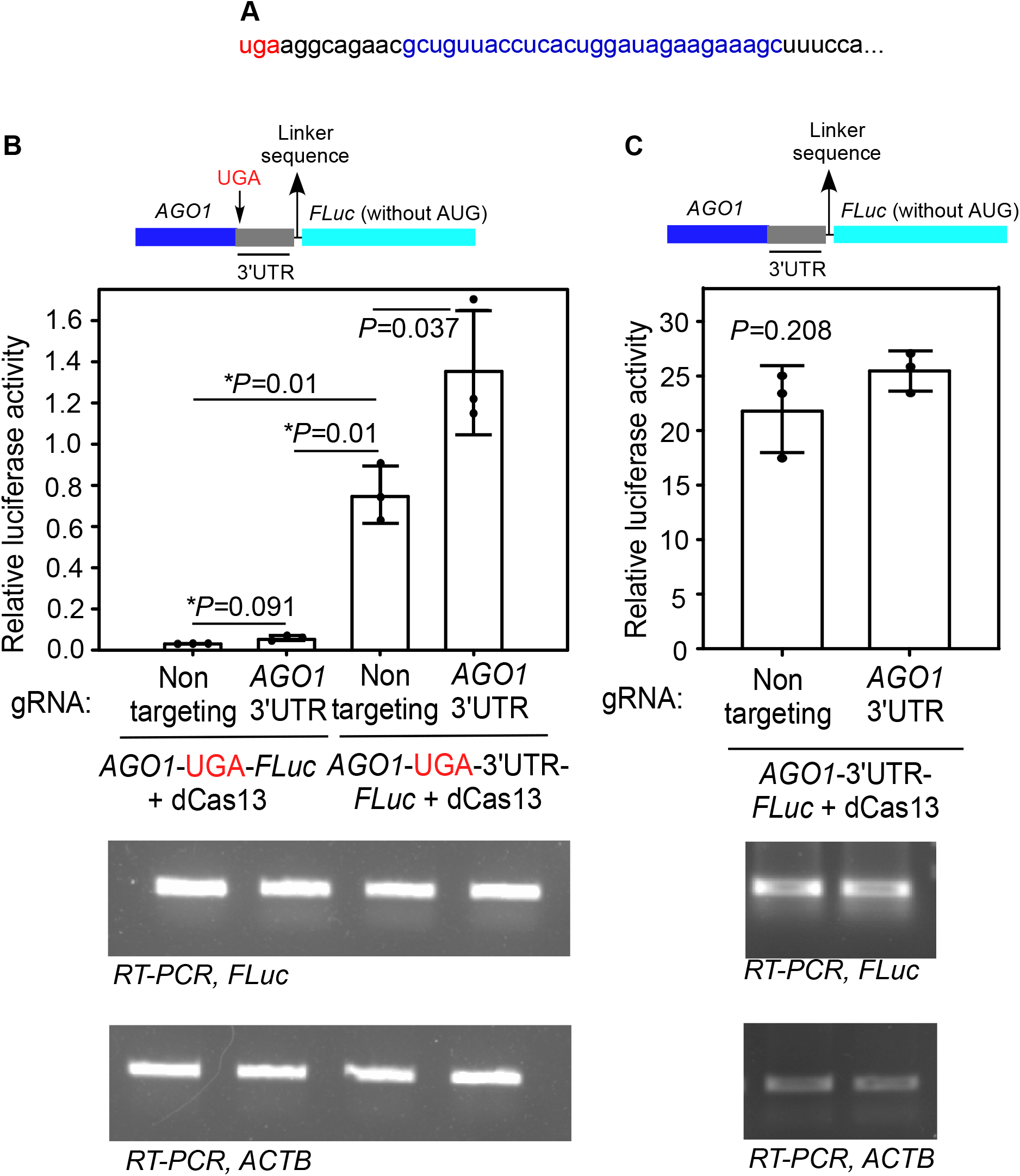
*AGO1*-3′UTR-targeting dCas13 enhances SCR of *AGO1* mRNA. **(A)** The sequence of the proximal 3′UTR of *AGO1*. The canonical stop codon (uga) and the gRNA targeting region are shown in red and blue, respectively. **(B)** Luminescence-based SCR assay. The cDNA of firefly luciferase (FLuc) was cloned downstream of and in frame with the partial cDNA of *AGO1* (696 nucleotides of the 3′ end) along with its proximal 3′UTR (99 nucleotides) such that luminescence is expected only after SCR (see schematic). The constructs were transfected in HEK293 cells along with plasmids expressing dCas13 and *AGO1*-3′UTR-targeting or non-targeting gRNA. The luminescence was measured 48 h after transfection. FLuc activity relative to the activity of the co-transfected renilla luciferase is shown. **(C)** Effect of CRISPR-dCas13 system on canonical translation and mRNA level. *AGO1*-3′UTR-*FLuc* construct without any stop codon in between was transfected in HEK293 cells along with plasmids expressing dCas13 and gRNA. Relative luciferase activity was measured as described above. RT-PCR results show the expression of *FLuc* mRNA. Graphs are representatives of three independent experiments. Bars indicate mean ± sd (n=3). Two-sided Student’s t-test was used to calculate the *P* values. * Welch’s correction was applied.

To test if dCas13 can enhance the SCR of *AGO1*, we employed a luminescence-based readthrough assay (Singh *et al*., 2019). Part of the *AGO1* coding sequence and the proximal part of its 3′UTR were cloned upstream of and in-frame with the firefly luciferase (*FLuc*) coding sequence. Expression of luciferase, and therefore luminescence, is expected only if there is SCR across the canonical stop codon of *AGO1* (schematic in Fig. 1B). This luciferase construct was transfected in HEK293 cells along with constructs expressing dCas13 and the gRNA. As reported previously, we observed SCR in *AGO1* across its canonical stop codon in the presence of the proximal part of its 3′UTR as indicated by the increased luminescence (1^st^ and 3^rd^ bar in Fig 1B) (Singh *et al*., 2019). There was a modest enhancement in the SCR in cells expressing dCas13 and *AGO1*-3′UTR-targeting gRNA compared to those expressing a non-targeting gRNA (3^rd^ and 4^th^ bar Fig 1B). Since luminescence is an indicator of SCR, this assay suggests that dCas13 along with *AGO1*-3′UTR-targeting gRNA can enhance the SCR across the canonical stop codon of *AGO1*. Importantly, the combination of *AGO1*-3′UTR-targeting gRNA and dCas13 failed to alter the activity of the luciferase construct that lacked the proximal part of *AGO1* 3′UTR, which is the target region of the gRNA (1^st^ and 2^nd^ bars in Fig. 1B). This shows that the gRNA and dCas13 work in a transcript-specific manner to enhance SCR. Furthermore, this combination did not show detectable change in the normal translation as indicated by the luminescence in cells transfected with *AGO1*-*3′UTR*-*FLuc* construct without any stop codon between them. Similarly, RT-PCR analyses revealed no change in the level of the target RNA, i.e, *AGO1*-luciferase mRNA in this case (Fig 1C). Similar observations were made in *MTCH2*, another mRNA known to undergo SCR (Eswarappa *et al*., 2014; Manjunath *et al*., 2020). dCas13 coupled with *MTCH2-*3′UTR-targeting gRNA enhanced SCR across the canonical stop codon of *MTCH2* in a transcript-selective manner without altering the levels of its mRNA or its canonical translation (Fig EV2). Together, these results show that CRISPR-dCas13 system can be used to enhance the SCR without affecting the target mRNA level.

### dCas13 enhances the stop codon readthrough of endogenous *AGO1*

We next tested if dCas13 can increase the stop codon readthrough of endogenous *AGO1,* which results in Ago1x. HEK293 cells transfected with plasmids expressing *AGO1*-3′UTR-targeting gRNA and dCas13 showed about 3-fold increase in the expression of Ago1x protein compared to those transfected with a non-targeting gRNA. However, there was no detectable change in the total Ago1 protein and *AGO1* mRNA levels under the same conditions (Fig 2A and B). The gRNA-mediated enhancement of Ago1x expression was observed using dCas13 from another bacterium, *Porphyromonas gulae* (dPguCas13) also (Fig 2C). These results show that the designed *AGO1*-3′UTR-targeting gRNA increases the expression of Ago1x, the readthrough product of *AGO1*. Importantly, this action does not affect the canonical translation of *AGO1* mRNA or its levels. This implies that the increased Ago1x expression is because of increased SCR across the canonical stop codon of *AGO1* mRNA. These observations are consistent with the luminescence-based readthrough assays described above (Fig 1). The increase in Ago1x level was achieved only when both dCas13 and *AGO1*-3′UTR-targeting guide RNA were expressed in cells; the gRNA alone did not enhance Ago1x expression (Fig EV3A). This shows that increased expression of Ago1x is the result of the gRNA-mediated action of dCas13.

**Figure 2.**
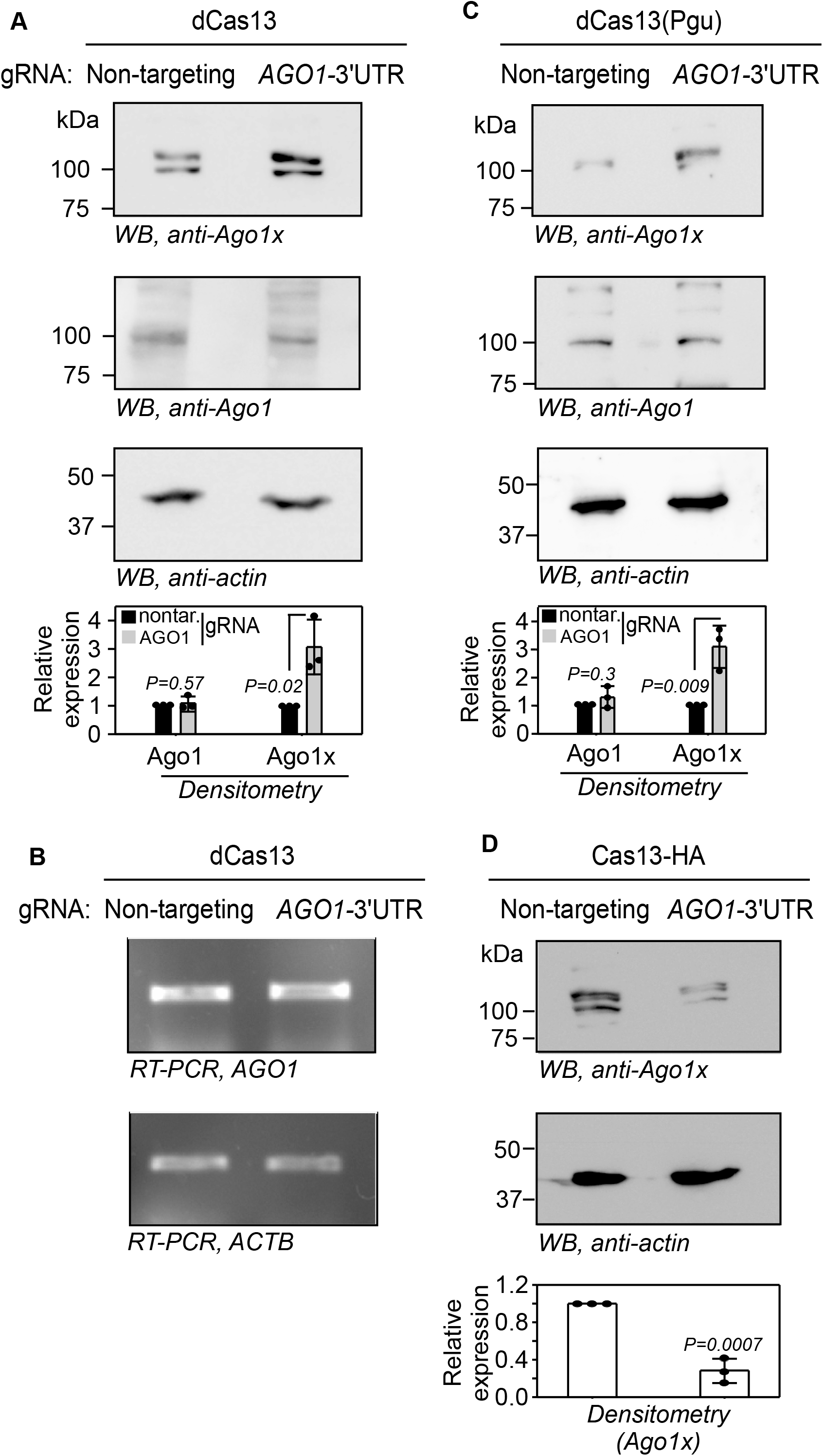
*AGO1*-3′UTR-targeting dCas13 enhances the expression of Ago1x, the SCR product of endogenous *AGO1* mRNA. **(A)** Western blot showing the expression of Ago1x and Ago1 in HEK293 cells expressing dCas13 (from *L. wadei*) and *AGO1*-3′UTR-targeting gRNA. **(B)** RT-PCR result showing the expression of *AGO1* mRNA in same cells. **(C)** Western blot showing the expression of Ago1x and Ago1 in HEK293 cells expressing dCas13 (from *P. gulae*) and *AGO1*-3′UTR-targeting gRNA. **(D)** Western blot showing reduced expression of Ago1x in HEK293 cells expressing Cas13a and *AGO1*-3′UTR-targeting gRNA. Graphs in all panels represent densitometry values (mean ± sd, N=3). Two-sided Student’s t-test was used to calculate the *P* values.

The same *AGO1*-3′UTR-targeting gRNA reduced the expression of Ago1x when transfected along with Cas13, which is catalytically active (Fig 2D). This result along with the results shown in Fig 2A and S3A show that the *AGO1*-3′UTR-targeting gRNA can guide Cas13 and dCas13 to its target mRNA, i.e., *AGO1*. Furthermore, targeting the same *AGO1*-3′UTR region (Fig 1A) on the genomic DNA using CRISPR-Cas9 system knocked down the expression of Ago1x (Fig EV3B). Complete knockout of Ago1x could not be achieved possibly because of lethality consistent with a previous report (Ghosh *et al*., 2020). This result shows that the band we observed in Western blot assay indeed represents the SCR product of *AGO1*, i.e., Ago1x.

Since overexpression of Ago1x can enhance global translation (Singh *et al*., 2019), we investigated the effect of dCas13-mediated induction of *AGO1* readthrough on global translation. For this we employed ribopuromycylation assay (Bastide *et al*, 2018). In this assay, puromycin is incorporated into nascent peptides during translation as it mimics tyrosyl-tRNA. This can be detected by Western blot using anti-puromycin antibody. We observed enhanced global translation in HEK293 cells transfected with dCas13 along with *AGO1*-3′UTR-targeting gRNA (Fig 3). This is consistent with the increased Ago1x expression in those cells. These results show that the induction of translational readthrough of *AGO1* by dCas13 results in a functional product.

**Figure 3.**
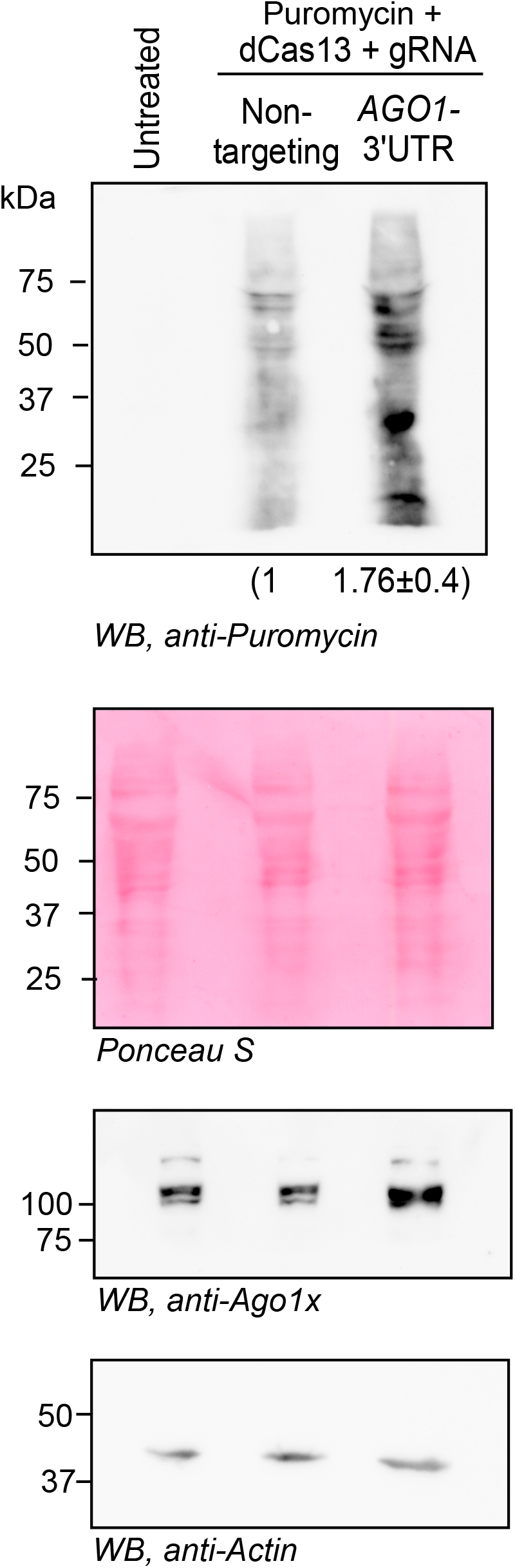
*AGO1*-3′UTR-targeting dCas13 enhances global translation. Ribopuromycylation assay was performed in HEK293 cells expressing dCas13 and *AGO1*-3′UTR-targeting gRNA as described in Methods. Numbers below indicate the densitometry values (normalized to Ponceau-S staining) (Mean ± sd, N=3, *P*=0.027).

### dCas13 causes ribosomal pausing on target mRNA

Ribosomal pausing at the stop codon is associated with stop codon readthrough and ribosomal frameshifting induced by *cis*- and *trans*-acting factors and also by small molecules(Annibaldis *et al*, 2020; Bao *et al*, 2020; Dever *et al*, 2018; Lashkevich *et al*, 2020; Lawson *et al*, 2021; Seidman *et al*, 2011; Sharma *et al*, 2021). Therefore, we tested if dCas13 can cause ribosome pausing at the target mRNA. We adopted a previously established dual fluorescence-based ribosomal stalling assay (Juszkiewicz & Hegde, 2017). In this assay, GFP and mCherry are expressed from a single mRNA. P2A sites inserted between them (schematic in Fig 4A) ensure that the two proteins are independent. Ribosomal pausing or stalling between them would result in reduced mCherry signal relative to the GFP signal. As reported earlier, the presence of a strong ribosomal stalling sequence (20 repeats of lysine codon, AAA) resulted in a significant reduction in the ratio of the mean fluorescence intensity of mCherry to that of GFP in transfected cells. To investigate the effect of dCas13 binding, we introduced the proximal 3ʹUTR of *AGO1* mRNA (99 nucleotides, target of gRNA-dCas13) between the coding sequences of GFP and mCherry (schematic in Fig 4A). The construct was transfected in HeLa cells along with plasmids expressing dCas13 and *AGO1*-3′UTR-targeting gRNA. The fluorescence intensities were quantified by flow cytometry 24 h after transfection. We observed a significant reduction in the ratio of the mean fluorescence intensity of mCherry to that of GFP in cells expressing dCas13 and *AGO1*-3′UTR-targeting gRNA. This observation suggests ribosomal pausing caused by dCas13 binding.

**Figure 4.**
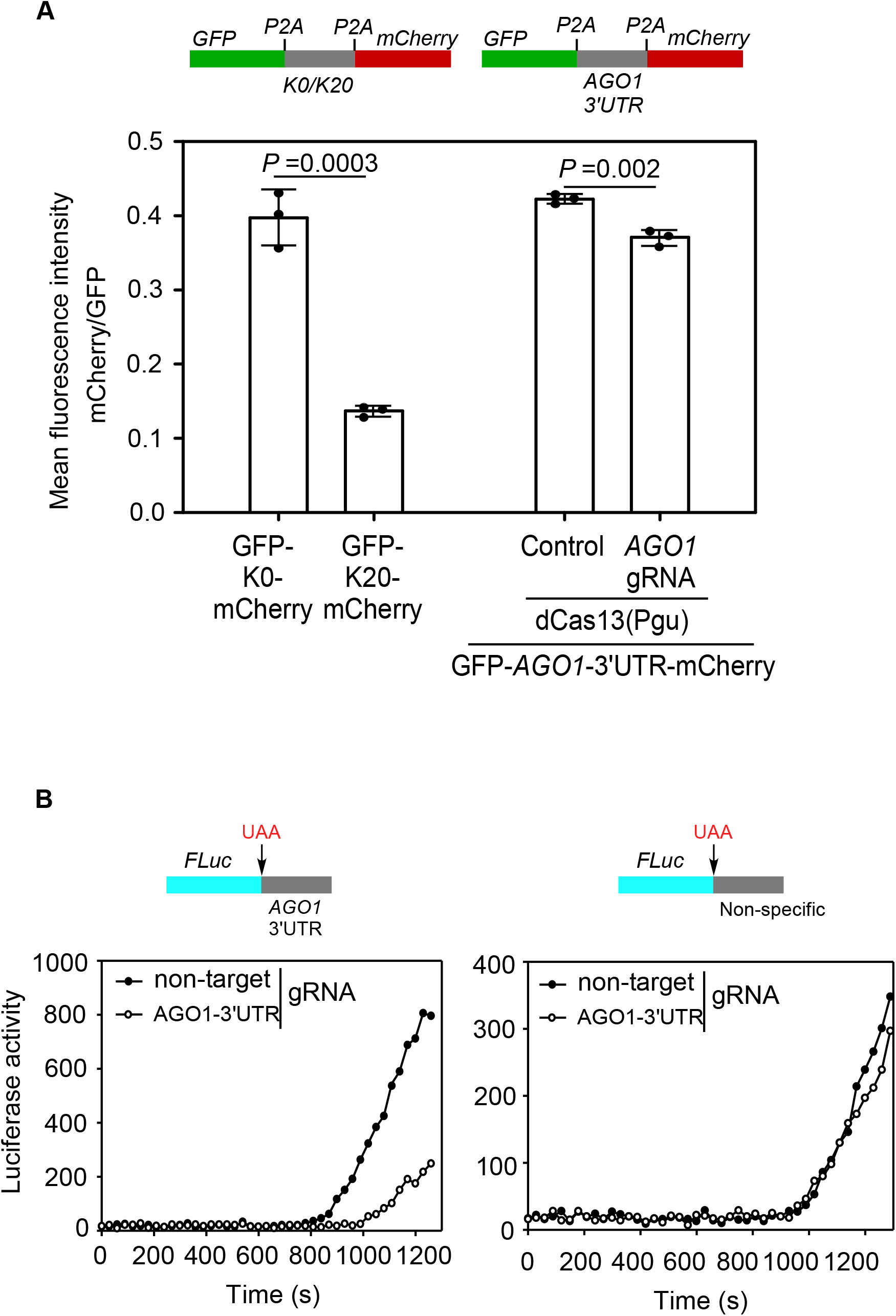
*AGO1*-3′UTR-targeting dCas13 causes ribosomal pausing. **(A)** Fluorescence-based ribosomal pausing assay. HeLa cells were transfected with a plasmid construct containing cDNAs of GFP and mCherry separated by proximal 3′UTR of *AGO1* (99 nucleotides). The graph shows the ratio of mean fluorescence intensity of mCherry to that of GFP in the presence of dCas13 (Pgu) and *AGO1*-3′UTR-targeting gRNA. GFP-mCherry constructs with 20 lysine codons (AAA) was used as positive control for the assay. Graph is a representative of three independent experiments. Bars show mean ± sd, n=3. Two-sided Student’s t-test was used to calculate the *P* values. **(B)** Luminescence-based ribosomal pausing assay. cDNA of FLuc was tagged with the proximal 3′UTR of *AGO1* (99 nucleotides) or a non-specific sequence of same length (see schematics). They were *in vitro* transcribed, and the resulting mRNA was *in vitro* translated using rabbit reticulocyte lysate in the presence of extracts from HEK293 cells expressing dCas13 (Pgu) and *AGO1*-3′UTR-targeting gRNA or non-targeting gRNA. FLuc activity (luminescence) was measured every 30 sec. Graphs are representatives of three independent experiments.

To further investigate ribosomal pausing, we used a luciferase-based assay. In this assay, the cDNA of firefly luciferase (FLuc) was tagged with the proximal 3′UTR of *AGO1* (99 nucleotides, target of gRNA-dCas13) at its 3′ end. Ribosomal pausing at the stop codon will cause a delay in the release of the newly synthesized luciferase protein and therefore its activity (i.e., luminescence), which can be quantified (Lashkevich *et al*., 2020). We performed *in vitro* transcription of these constructs. The obtained mRNAs were used for *in vitro* translation using rabbit reticulocyte lysate in the presence of extracts of HEK293 cells expressing dCas13 and *AGO1*-3′UTR-gRNA or non-targeting gRNA. We observed a delay in the appearance of the luminescence in the presence of *AGO1* gRNA compared to non-targeting gRNA suggesting ribosomal pausing. Importantly, this delay was not observed when FLuc was tagged with a non-specific sequence of same length (Fig 4B). Together, these two assays show that dCas13, like other readthrough-inducing *trans*-acting factors can cause ribosomal pausing. This pausing might provide sufficient opportunity for the near-cognate tRNAs to recognize the stop codon enabling readthrough.

### dCas13 enhances the stop codon readthrough of *VEGFA*

Next, we tested the ability of dCas13 to induce readthrough in another mRNA. We chose *VEGFA* mRNA, which encodes a secretory pro-angiogenic protein VEGF-A. The SCR of *VEGFA* results in a longer isoform termed VEGF-Ax with a unique C-terminus, which prevents its binding to Neuropilin 1, an important co-receptor in VEGF-A signalling. Because of this, VEGF-Ax shows anti-angiogenic or weakly pro-angiogenic properties (Eswarappa *et al*., 2014; Xin *et al*, 2016).

We designed a gRNA to target the *VEGFA* mRNA downstream of the canonical stop codon (Fig 5A). This gRNA was expressed in HEK293 cells along with dCas13, and the VEGF-Ax was detected in the conditioned medium using Western blot. The level of secreted endogenous VEGF-Ax was increased in cells transfected with *VEGFA*-3′UTR-targeting gRNA compared to those expressing non-targeting gRNA. However, there was no change in the level of *VEGFA* mRNA under same conditions (Fig 5B and 5C). These results show that dCas13, guided by the specific gRNA, enhances the expression of VEGF-Ax, the SCR product of *VEGFA*. The same *VEGFA*-3′UTR-targeting gRNA caused knockdown of VEGF-Ax when expressed along with Cas13 (catalytically active form) in these cells confirming the ability of this gRNA to target *VEGFA* mRNA (Fig 5D). Using CRISPR-Cas9 system, we generated VEGF-Ax knockout cells by targeting the proximal 3′UTR of *VEGFA* mRNA (Fig 5E), the region responsible for SCR. These cells showed complete absence of VEGF-Ax confirming that the 20 kDa band we observe in Western blots is the product of SCR of *VEGFA* (Fig 5E).

**Figure 5.**
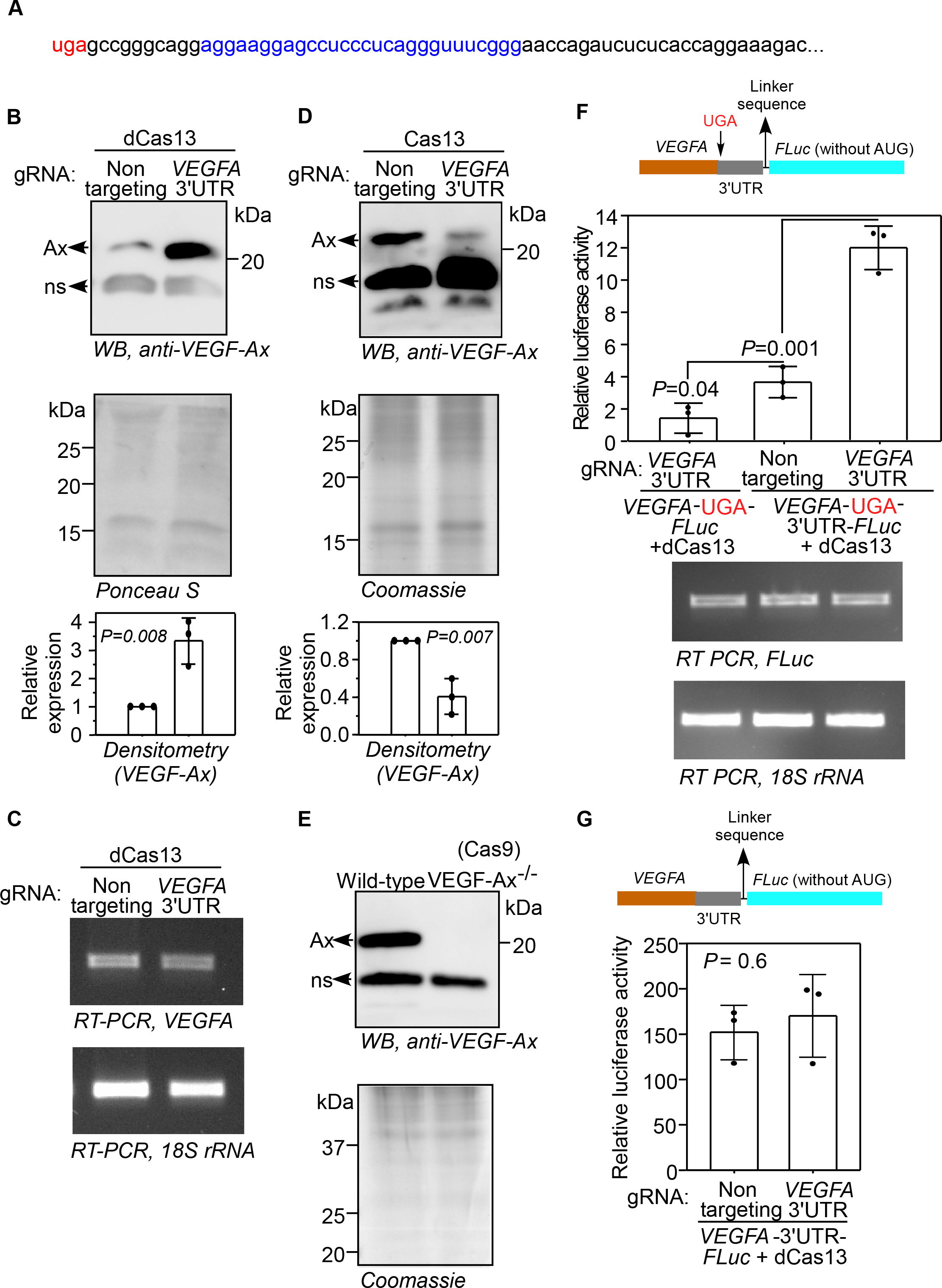
Enhancement of SCR of *VEGFA* using dCas13. **(A)** The sequence of the proximal 3′UTR of *VEGFA*. The region targeted by the gRNA is shown in blue. The canonical stop codon (uga) is shown in red. **(B)** Western blot showing the effect of dCas13 on the SCR of *VEGFA*. HEK293 cells were transfected with plasmids expressing dCas13 and *VEGFA*-3′UTR-targeting gRNA or non-targeting gRNA. The conditioned medium was collected 48 h after transfection and subjected to Western blot to detect VEGF-Ax (the SCR product of *VEGFA*) expression. **(C)** RT-PCR result showing the expression of *VEGFA* mRNA in those cells. **(D)** Western blot showing the level of VEGF-Ax in the conditioned medium of HEK293 cells expressing Cas13 and *VEGFA*-3′UTR-targeting gRNA. The conditioned medium was collected 48 h after transfection. **(E)** Western blot showing the absence of VEGF-Ax in the conditioned medium of VEGF-Ax knockout cells generated using CRISPR-Cas9 system. Graphs in (B) and (D) show densitometry values (mean ± sd, N=3). Ax, VEGF-Ax; ns, non-specific band. **(F)** Luminescence-based SCR assay. The cDNA of firefly luciferase (FLuc) was cloned downstream of and in frame with the cDNA of *VEGFA a*long with its proximal 3′UTR (63 nucleotides) such that luminescence is expected only after SCR (see schematic). The constructs were transfected in HEK293 cells along with plasmids expressing dCas13 and *VEGFA*-3′UTR-targeting or non-targeting gRNA. The luminescence was measured 24 h after transfection. FLuc activity relative to the activity of the co-transfected renilla luciferase is shown. Bottom panel shows the expression of *FLuc* mRNA. **(G)** Effect of dCas13 on canonical translation. *VEGFA*-3′UTR-FLuc construct without any stop codon in between was transfected in cells along with plasmids expressing dCas13 and gRNAs. Relative luciferase activity was measured as described above. Graphs in (F) and (G) are representative of three independent experiments. Bars indicate mean ± sd (n=3). Two-sided Student’s t-test was used to calculate the *P* value.

Next, luminescence-based readthrough assay was performed to investigate the ability of dCas13 to enhance the SCR of *VEGFA*. *VEGFA* coding sequence and the proximal part of its 3′UTR were cloned upstream of and in-frame with firefly luciferase coding sequence (schematic in Fig. 5F) (Eswarappa *et al*., 2014). This construct was expressed in HEK293 cells along with dCas13 and the gRNA. SCR across the stop codon results in luciferase expression which can be quantified as luminescence. We observed increased luminescence in cells expressing *VEGFA*-3′UTR-targeting gRNA compared to those expressing non-targeting gRNA (2^nd^ and 3^rd^ bars in Fig 5F) showing enhancement of SCR. There was no change in the luciferase mRNA level in any of these conditions (Fig 5F). Furthermore, the translation of *VEGFA*-3′UTR-luciferase construct, which did not have any stop codon in between, was unaltered by the expression of *VEGFA*-3′UTR-targeting gRNA and dCas13 (Fig 5G). Together, these results show that CRISPR-dCas13 system can be used to enhance SCR across the canonical stop codon of *VEGFA*, without affecting the cellular levels of *VEGFA* mRNA.

Because VEGF-Ax is anti-angiogenic (Eswarappa *et al*., 2014) or weakly angiogenic (Xin *et al*., 2016), compared to the canonical isoform VEGF-A, SCR of *VEGFA* mRNA will result in a net anti-angiogenic effect. Therefore, enhancement of SCR in *VEGFA* will be useful to treat diseases with excessive and abnormal angiogenesis such as cancer and retinopathies.

### dCas13 can be programmed to induce translational readthrough across premature termination codons (PTC)

Nonsense mutations resulting in PTCs are responsible for about 11% of genetic diseases. We investigated if dCas13 can be used to induce translational readthrough across PTCs. For this we used GFP construct with a PTC at 57^th^ codon (UGG to UGA or W57* mutation). We designed a gRNA that targets the region downstream of the PTC. In HEK293 cells we observed full-length GFP in cells expressing dCas13 along with the GFP-targeting gRNA, but not in cells expressing non-targeting gRNA, showing readthrough across the PTC (Fig 6A).

**Figure 6.**
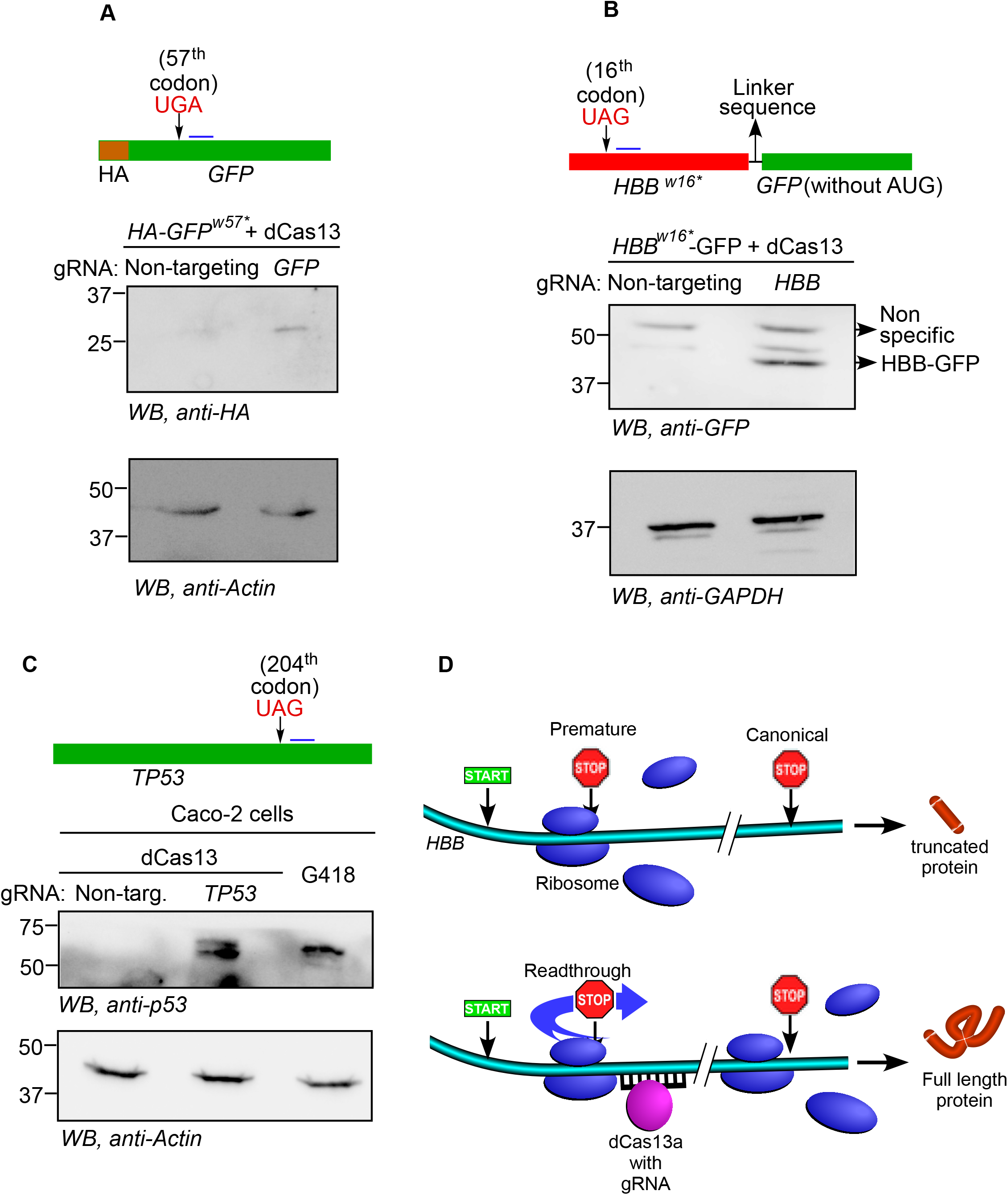
Induction of SCR across premature termination codons (PTC) using dCas13. **(A)** Western blot showing the expression of full-length HA-tagged GFP from a construct with PTC at the 57^th^ codon of GFP coding sequence. The construct was transfected in HEK293 cells along with plasmids expressing dCas13 and a GFP-specific gRNA or a non-targeting gRNA. **(B)** Western blot showing the expression of full-length GFP-tagged human β-globin from a construct with PTC at the 16^th^ codon of *HBB*. The construct was transfected in HEK293 cells along with plasmids expressing dCas13 and a *HBB*-specific gRNA or a non-targeting gRNA. **(C)** Western blot showing the expression of full-length human p53 in Caco-2 cells whose genome has a PTC at 204^th^ codon of *TP53* gene. Cells were transfected with plasmids expressing dCas13 and a *TP53*-specific gRNA or a non-targeting gRNA. G418, a known readthrough-inducing agent was used as a positive control. All results are representatives of three independent experiments. Schematics indicate the approximate position of the PTCs and the gRNA target region (horizontal blue lines next to PTCs). **(D)** Schematic showing the induction of SCR across PTCs by dCas13.

β-thalassemia is a condition caused by nonsense mutations in *HBB* gene (encodes β-globin protein). This condition is characterized by reduced haemoglobin level. Worldwide, 80 to 90 million people carry thalassemia allele in *HBB* gene (Origa, 2017). Any strategy that can induce SCR across the premature stop codons can provide therapeutic benefit to β-thalassemia patients with nonsense mutations. We tested the ability of CRISPR-dCas13 system to induce SCR in β-thalassemia context. We used an *HBB* construct with a PTC at its 16^th^ codon (TGG to TAG or W16* mutation) as described previously (Fig. 6B) (Kar *et al*., 2020). This nonsense mutation (w16*) is frequently found in β-thalassemia patients (Colah *et al*, 2009; Kazazian *et al*, 1984). The coding sequence of GFP was cloned in-frame with and downstream of *HBB^w16*^* (schematic in Fig. 6B) such that readthrough across the thalassemia-causing PTC will result in full-length β-globin protein tagged with GFP. This construct along with plasmids expressing dCas13 and *HBB*-targeting gRNA (targets downstream of the PTC) were transfected in HEK293 cells. While β-globin was not detected in cells expressing non-targeting gRNA, we could detect full length β-globin-GFP protein in cells expressing *HBB*-targeting gRNA (Fig. 6B). These results show that dCas13 can be used to induce SCR across the thalassemia-causing PTC in *HBB* mRNA.

Next, we used Caco-2 cells (human colorectal adenocarcinoma cell line) which harbour a nonsense mutation in *TP53* gene (GAG to TAG or E204* mutation). These cells do not express p53 protein because of this mutation. We observed p53 expression in these cells when they were transfected with plasmids expressing dCas13 and a *TP53*-targeting gRNA (targets downstream of the PTC). The p53 expression was comparable to that induced by G418 (Geneticin), an aminoglycoside known to induce SCR (Fig 6C).

Together, these results demonstrate the ability of dCas13 to induce SCR across PTCs. This property can be utilized in the treatment of diseases caused by nonsense mutations such as β-thalassemia, Duchenne muscular dystrophy, haemophilia, etc. Since gRNAs ensure that dCas13 targets a specific mRNA, this strategy can achieve high specificity unlike other SCR-inducing agents such as ataluren and aminoglycoside. Also, this system can be personalized to a patient depending on the location of the nonsense mutation.

Overall, using multiple mRNAs – *AGO1*, *VEGFA, MTCH2*, *HBB, GFP and TP53* – we have demonstrated that dCas13 can be programmed with the help of a gRNA to enhance (or induce) SCR across the canonical (or premature) stop codons. Importantly, this is achieved in a transcript-selective and stop codon-specific manner without altering the transcript level. This specificity provides a key advantage over the existing SCR-inducing strategies/molecules, which are largely nonselective. Thus, induction of SCR is a new addition to the CRISPR system’s expanding arsenal of biotechnological applications directed towards therapeutics.

## Materials and Methods

(Reagents table is uploaded as a separate file)

### Cell culture

HEK293, HeLa, Caco-2 cells were cultured using Dulbecco’s Modified Eagle’s Medium (DMEM, HiMedia), which was supplemented with 10% fetal bovine serum (FBS, Gibco) and 1% antibiotics (10000 U/ml penicillin, 10000 µg/ml streptomycin, Lonza). The cells were incubated in a humidified atmosphere at 37°C with 5% CO_2_.

### CRISPR-Cas13 system: Construction of plasmids

The reporter (Luciferase/GFP) constructs (in pcDNA3.1B vector background) used for SCR assays for *AGO1*, *VEGFA, MTCH2* and *HBB* have been described previously (Eswarappa *et al*., 2014; Kar *et al*., 2020; Manjunath *et al*., 2020; Singh *et al*., 2019). In case of the GFP reporter for SCR across a premature termination codon, the GFP sequence with a nonsense mutation at the 57^th^ codon was amplified from pMHG-W57* reporter, a gift from Daniel Liebler (Addgene plasmid #72850) (Halvey *et al*, 2012). It was cloned in pIRESneo-FLAG/HA plasmid to express the reporter with an N-terminal FLAG-HA tag.

The GFP-K0-mCherry and GFP-K20-mCherry reporter constructs for ribosomal stalling were a kind gift from Prof. Ramanujan Hegde, MRC Laboratory of Molecular Biology, Cambridge (Juszkiewicz & Hegde, 2017). The proximal 3ʹUTR of *AGO1* mRNA was cloned between the GFP and mCherry reporters. For the luciferase-based ribosomal pausing assay, the proximal 3′UTR of *AGO1* mRNA was cloned downstream of the coding region of *FLuc,* separated by a stop codon (in pcDNA3.1B vector backbone). As a length-matched control, a non-specific sequence was cloned in place of the *AGO1* proximal 3′UTR.

pC014-LwCas13a-msfGFP plasmid expressing active Cas13, pC015-dLwCas13a-NF expressing the catalytically inactive mutant of Cas13 (dCas13) and pC016-LwCas13a guide expression backbone (with U6 promoter) were a gift from Feng Zhang (Addgene plasmids #91902, #91905 and #91906)(Abudayyeh *et al*., 2017). pHAGE-IRES-puro-NLS-dPguCas13b-EGFP-NLS-3xFlag vector, which expresses dCas13 from *Porphyromonas gulae*, was a gift from Ling-Ling Chen (Addgene plasmid # 132400)(Yang *et al*., 2019). pLentiRNACRISPR_001-hU6-DR_BsmBI-EFS-PguCas13b-NLS-2A-Puro -WPRE expressing PguCas13b was a gift from Neville Sanjana (Addgene plasmid # 138143) (Wessels *et al*, 2020). This was modified in this study to express dPguCas13 instead of PguCas13. Oligos were cloned in the guide expression backbones to generate either the 28-nucleotide or 30-nucleotide gRNAs targeting the desired region.

The gRNA sequences are given below (5′ to 3′): *AGO1*: GCTTTCTTCTATCCAGTGAGGTAACAGC/ AAGCTTTCTTCTATCCAGTGAGGTAACAGC *MTCH2:* CTGGGACTACAGAAATGTCACTGTCCCT *VEGFA*: CCCGAAACCCTGAGGGAGGCTCCTTCCT *HBB* (W16*): GGGCCTCACCACCAACTTCATCCACGTT *GFP* (W57*) – CACCATAAGTCAGGGTGGTCACCAGAGT *TP53* (E204*) – CCACACTATGTCGAAAAGTGTTTCTGTC Non-targeting (Abudayyeh *et al*., 2017): TAGATTGCTGTTCTACCAAGTAATCCAT

### CRISPR-Cas9 mediated deletion

sgRNAs (sequences given below) were cloned in pSpCas9 (BB)-2A-Puro plasmid. Cells were transfected with these plasmids (two per gene) using Lipofectamine 2000 (Thermo Fisher Scientific). 24 h post-transfection, transfected cells were selected using 2 μg/ml of puromycin (Sigma) for 5 days. The surviving cells were reseeded in a 96-well plate at a density of single cell per well. These clones were expanded and screened for the required genetic deletion by PCR. The deletion was confirmed by sequencing of the PCR product and by Western blot.

sgRNA sequences (5′ to 3′):

*AGO1*: GCAGAACGCTGTTACCTCAC and GCTGTGCCACCCAAATCCAG

*VEGFA:* ACAAGCCGAGGCGGTGAGCC and GGAAAGACTGATACAGAACG

### Luminescence-based translational readthrough assays

HEK293 cells were seeded in 24-well plates at 70%-80% confluency. Conditions were optimized to achieve >75% transfection efficiency using pEGFP-C1. 200 ng/well of firefly luciferase-encoding reporter plasmids, 500 ng/well each of LwCas13a or dLwCas13a-NF and gRNA (gene-specific or non-targeting) were transfected using Lipofectamine 2000. 10 ng/well of *Renilla* luciferase was used as transfection control. Firefly and *Renilla* luciferase activities were measured using Dual-Luciferase Reporter Assay System (Promega) using GloMax Explorer (Promega) 24 h post-transfection in case of samples involving *VEGFA and MTCH2*, and 48 h post-transfection in case of *AGO1*.

### Antibodies

Antibodies specific to the readthrough region of *VEGFA* (against the peptide AGLEEGASLRVSGTR) and *AGO1* (against the peptide RQNAVTSLDRRKLSKP) were generated as described in previous studies (Eswarappa *et al*., 2014; Singh *et al*., 2019). Anti-Ago1 antibody (Novus Biologicals, NB100-2817), Anti-GFP antibody (BioLegend, 902602), anti-VEGFA antibody (Thermo Fisher JH121, MA5-13182), anti-puromycin antibody (PMY-2A4, Developmental Studies Hybridoma Bank), anti-p53 antibody (AHO0152, Thermo Fisher), anti-FLAG antibody (Sigma, F1804), anti-GAPDH antibody (Sigma, G9295), anti-HA antibody (Sigma, 11867423001), anti-Actin antibody (Sigma, A3854) and horseradish-peroxidase-conjugated secondary antibodies (Thermo Fisher) were used as per the manufacturer’s instructions.

### Western blot-based stop codon readthrough assays

HEK293 cells were seeded in 6-well plates at 70%-80% confluency. For assays involving Cas13a-mediated knockdown of endogenous proteins, 2 µg/well of LwCas13a-NF along with 2 µg/well of gRNA-expressing plasmid (gene-specific or non-targeting) were transfected using Lipofectamine 2000. 24 h post-transfection the cell pellets (for Ago1x) or 48 h post-transfection the conditioned medium (for VEGF-Ax) was harvested for Western blotting. For dCas13-mediated induction of endogenous readthrough proteins, either 3 µg/well of dLwCas13a-NF or 3 µg/well of dPguCas13b-3xFLAG, along with 2 µg/well of gRNA-expressing plasmid (gene-specific or non-targeting) were transfected using Lipofectamine 2000. 24 h post-transfection the cell pellets (for Ago1 and Ago1x) or 48 h post-transfection the conditioned medium (for VEGF-Ax) was harvested for Western blotting. The conditioned medium was subjected to trichloroaceticacid (TCA) precipitation.

For the assays involving exogenous plasmids with PTCs, 3 µg/well of FLAG-HA-GFPw57*, 3 µg/well of dLwCas13a-NF along with 2 µg/well of gRNA-expressing plasmid (gene-specific or non-targeting) were transfected in HEK293 cells using Lipofectamine 2000. For the assay involving the premature stop codon in *HBB* mRNA, 500 ng/well of the GFP-encoding construct (*HBB^w16*^-GFP)*, 1 µg/well each of dLwCas13a-NF and gRNA-expressing plasmid (gene-specific or non-targeting) were transfected in HEK293 cells using Lipofectamine 2000. 24 h post-transfection, the cell pellets were harvested.

For assays involving SCR across endogenous premature stop codon, Caco-2 cells were seeded in 6-well plates at 80-90% confluency. They were transfected with 4 µg/well of dLwCas13a-NF along with 4 µg/well of gRNA-expressing plasmid (gene-specific or non-targeting) using Lipofectamine 2000. 48 h post-transfection, the cell pellets were harvested.

Cell pellets were lysed in cell lysis buffer (20 mM Tris-HCl, 150 mM NaCl, 1 mM EDTA, 1% Triton-X with protease inhibitor cocktail (Promega)) and subjected to Western blotting. Protein Assay Dye Reagent (Bio-Rad) was used to determine the protein concentration. 50-100 µg of the cell lysate was subjected to denaturing SDS-PAGE in an 8% or 10% or 12.5% or 15% Tris-glycine gel. After the transfer of proteins onto a PVDF membrane (Merck), blocking was carried out (5% skimmed milk in PBS). Following this, the membrane was probed with the specific primary antibody and then with the respective horseradish peroxidase-conjugated secondary antibody. Clarity ECL reagent (Bio-Rad) was used for the development of the blots, and the images were recorded using LAS-4000 imager (Fujifilm) or ChemiDoc Imaging System (Bio-Rad). Band intensities were quantified using ImageJ.

### RNA isolation and RT-PCR

RNA isolation was carried out using RNAiso Plus (TaKaRa). cDNA synthesis was carried out with 1 µg of RNA using oligo(dT) primers or gene-specific reverse primer and RevertAid Reverse Transcriptase (Thermo Fisher). Semi-quantitative analysis of the mRNA levels was carried out using gene-specific primers.

Primer sequences are as shown below (5ʹ to 3ʹ):

*FLuc*: CAACTGCATAAGGCTATGAAGAGA; ATTTGTATTCAGCCCATATCGTTT *ACTB* (β-Actin): AGAGCTACGAGCTGCCTGAC; AGCACTGTGTTGGCGTACAG *GFP*: ATGGTGAGCAAGGGCGAGGAGCTG; CTTGTACAGCTCGTCCATGCCGAG *AGO1:* GGGAGCCACATATCGGGGCAG; CTACCCCACCTCCCTCCTCCTTG 18s *rRNA:* GGCCCTGTAATTGGAATGAGTC; CCAAGATCCAACTACGAGCTT *VEGFA:* CTTGCCTTGCTGCTCTACC; CACACAGGATGGCTTGAAG

### Ribopuromycylation

HEK293 cells were seeded in 6-well plates at 70%-80% confluency. 3 µg/well of dLwCas13a-NF along with 2 µg/well of gRNA-expressing (*AGO1*-3′UTR-specific or non-targeting) plasmids were transfected using Lipofectamine 2000. 24 h post-transfection, cells were treated with 91 µM puromycin (Sigma) for 10-15 minutes at 37°C. Following puromycin treatment, cells were lysed and 50 µg of the cell lysate was subjected to Western blotting using anti-puromycin antibody. Ponceau S images of the PVDF membranes was used for normalization for densitometry analysis.

### Dual fluorescence-based ribosomal pausing assay

HeLa cells were seeded in 24-well plates at 70-80% confluence. 200 ng/well of the fluorescence reporter constructs along with 1 µg/well of dPguCas13b-gRNA construct were transfected using Lipofectamine 2000 following the manufacturer’s protocol. 24 h post-transfection, the samples were subjected to flow cytometry analysis using Cytoflex LX (Beckman Coulter).

### Luminescence-based ribosomal pausing assay

HEK293 cells were seeded in 6-well plates at 70-80% confluency. 2 µg/well each of plasmids expressing dPguCas13 and gRNA (either non-specific or gene-specific) were transfected using Lipofectamine 2000. 48 h post-transfection, the cells were resuspended in hypotonic extraction buffer (20mM HEPES (pH 7.5), 10mM Potassium acetate, 1mM Magnesium chloride and 4mM DTT), homogenized at 3000 rpm using a motor-driven tissue grinder (Genetix), and kept on ice for 15 min. Samples were centrifuged at 15000 rpm, for 20 min at 4° C and the supernatant was taken as the cell extract. The luciferase reporter constructs were linearized using *Xho*I and *in vitro* transcribed using T7 RNA polymerase (Thermo Scientific). 250 ng of reporter RNA was *in vitro* translated using rabbit reticulocyte lysate in the presence of 4 µg/µl cell extract, 0.5 mM D-Luciferin, and RNase inhibitor. Luminescence was measured every 30 s using the GloMax Explorer (Promega).

### RNA Immunoprecipitation

HEK293 cells were seeded in 6-well plates at 70%-80% confluency. Transfection was carried out with 3 µg/well of dPguCas13b-3xFLAG along with 2 µg/well of gRNA-expressing (non-targeting or gene specific) plasmids. 48 h post-transfection, the cells were washed with ice-cold PBS and fixed using 0.2% paraformaldehyde. After 15 min, the cells were treated with 125 mM glycine for 10 min to quench the crosslinking. The cells were washed with ice-cold PBS and lysed using RIPA lysis buffer with RNase inhibitor and Protease inhibitor cocktail. Cell lysates were incubated with anti-FLAG M2 Affinity Gel (SIGMA, A2220) overnight with tumbling at 4°C. Laemmli buffer was used to extract the immunoprecipitated proteins, which were subjected to Western blotting. Immunoprecipitated RNA was extracted using TaKaRa RNAiso Plus. cDNA synthesis was carried out using oligo(dT) reverse primer. Quantitative real-time was carried out using TB Green Premix Ex TaqII (TaKaRa) in CFX96 real-time PCR system (Bio-Rad). The PCR conditions: 95°C for 5 min, 40 cycles of 95°C for 30 sec, 56°C for 30 s, and 72°C for 30 s, followed by a single final extension step at 72°C for 5 min. Enrichment of *AGO1* mRNA in IP samples relative to the input samples was calculated using the 2^-ΔΔCt^ method. Sequences (5′ to 3′) of primers are as follows:

*ACTB* (β-Actin): AGAGCTACGAGCTGCCTGAC; AGCACTGTGTTGGCGTACAG

*AGO1:* GGGAGCCACATATCGGGGCAG; CTACCCCACCTCCCTCCTCCTTG

### Statistics

Two-sided Student’s t-test was used to test for the significance of the differences observed between samples in the experiments when samples showed normal distribution.

## Acknowledgements

This work was supported by the Department of Biotechnology (DBT), India (grant no. BT/PR38405/GET/119/309/2020). SME is a recipient of the Swarnajayanti Fellowship (SB/SJF/2020-21/18) from the Department of Science and Technology (DST)-Science and Engineering Research Board (SERB), India. Authors gratefully acknowledge the financial support from STARS grant from the Ministry of Education, DST Funds for Improvement of S&T infrastructure, and funds from the University Grants Commission, India. LEM acknowledges the Research Associateship received from the Indian Council for Medical Research.

## Author contributions

SME and LEM conceived the project and designed the experiments. LEM performed most of the experiments. AS and DK generated constructs used in SCR assays. AS and KS generated knockout cell lines. SME obtained funds and supervised the project. SME and LEM analysed the data and wrote the first draft. The manuscript was read and approved by all authors.

## Legends to Expanded View Figures

**Figure EV1.**
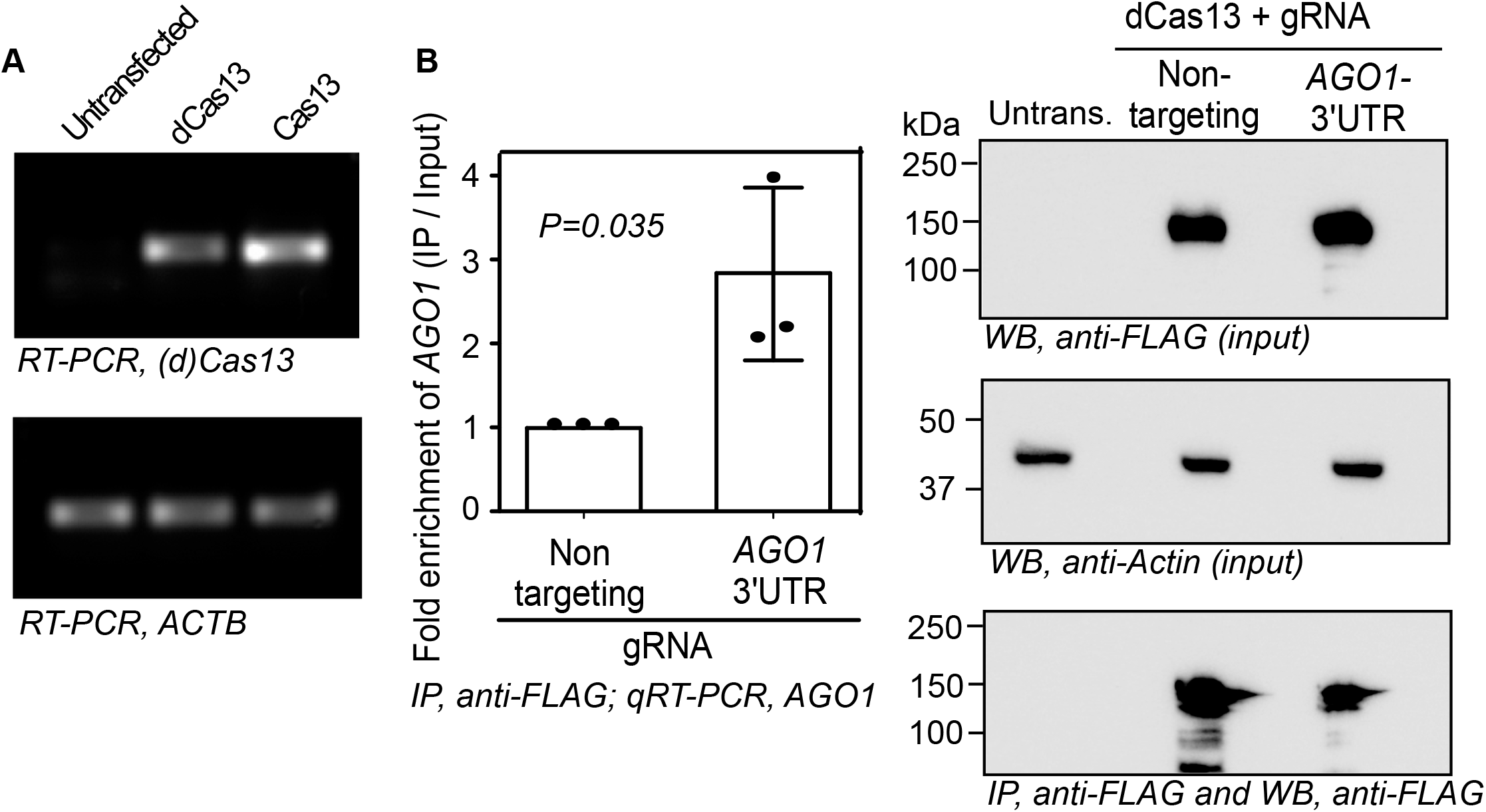
(A) RT-PCR analysis showing the expression of Cas13 and dCas13 (both from *Leptotrichia wadei*) in transfected HEK293 cells. (B) dCas13 interacts with its target mRNA, *AGO1*, mediated by a specific gRNA. Constructs expressing dPguCas13b-3xFLAG along with gRNAs were transfected in HEK293 cells. Cell lysates were subjected to immunoprecipitation followed by RNA isolation and qRT-PCR to detect the enrichment of *AGO1* mRNA. Immunoprecipitates were also used for Western blot. Results are representative of three independent experiments (Graph, mean ± sd).

**Figure EV2.**
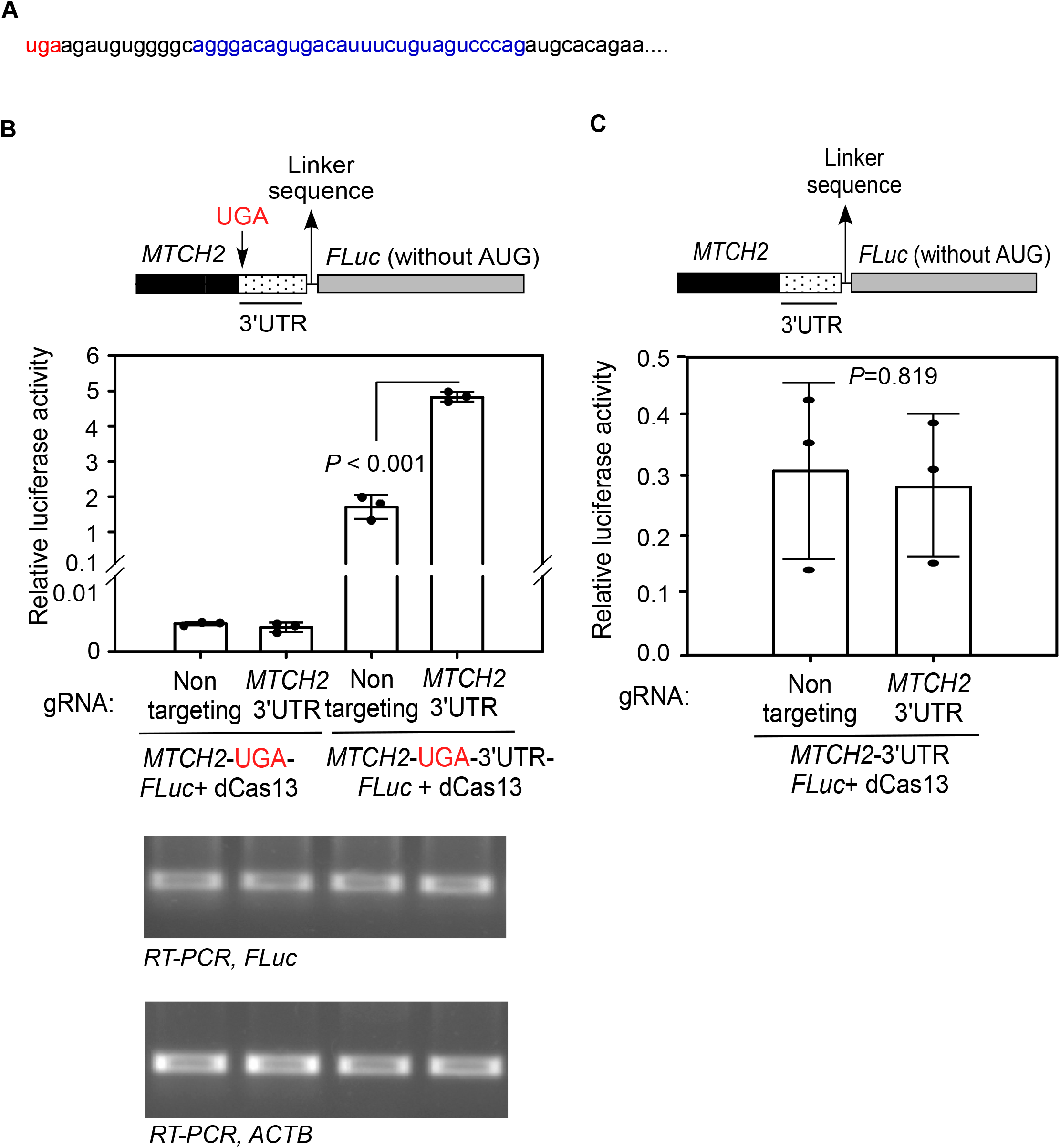
Enhancement of SCR across the canonical stop codon of *MTCH2* using CRISPR-dCas13 system. (A) The sequence of the proximal 3′UTR of *MTCH2*. The canonical stop codon (UGA) and the gRNA targeting region are in red and blue, respectively. (B) Luminescence-based SCR assay. The indicated constructs were transfected in HEK293 cells and firefly luciferase (FLuc) activity was measured 24 h after transfection. FLuc activity relative to the activity of the co-transfected renilla luciferase is shown. Bottom panel shows the expression of *FLuc* mRNA. (C) Effect of CRISPR-dCas13 system on normal translation. *MTCH2*-3′UTR-FLuc construct without any stop codon in between was transfected in cells along with plasmids expressing dCas13 and indicated gRNA. Relative luciferase activity was measured as described above. Graphs are representatives of three independent experiments. Bars indicate mean ± sd. Two-sided Student’s t-test was used to calculate the *P* value.

**Figure EV3.**
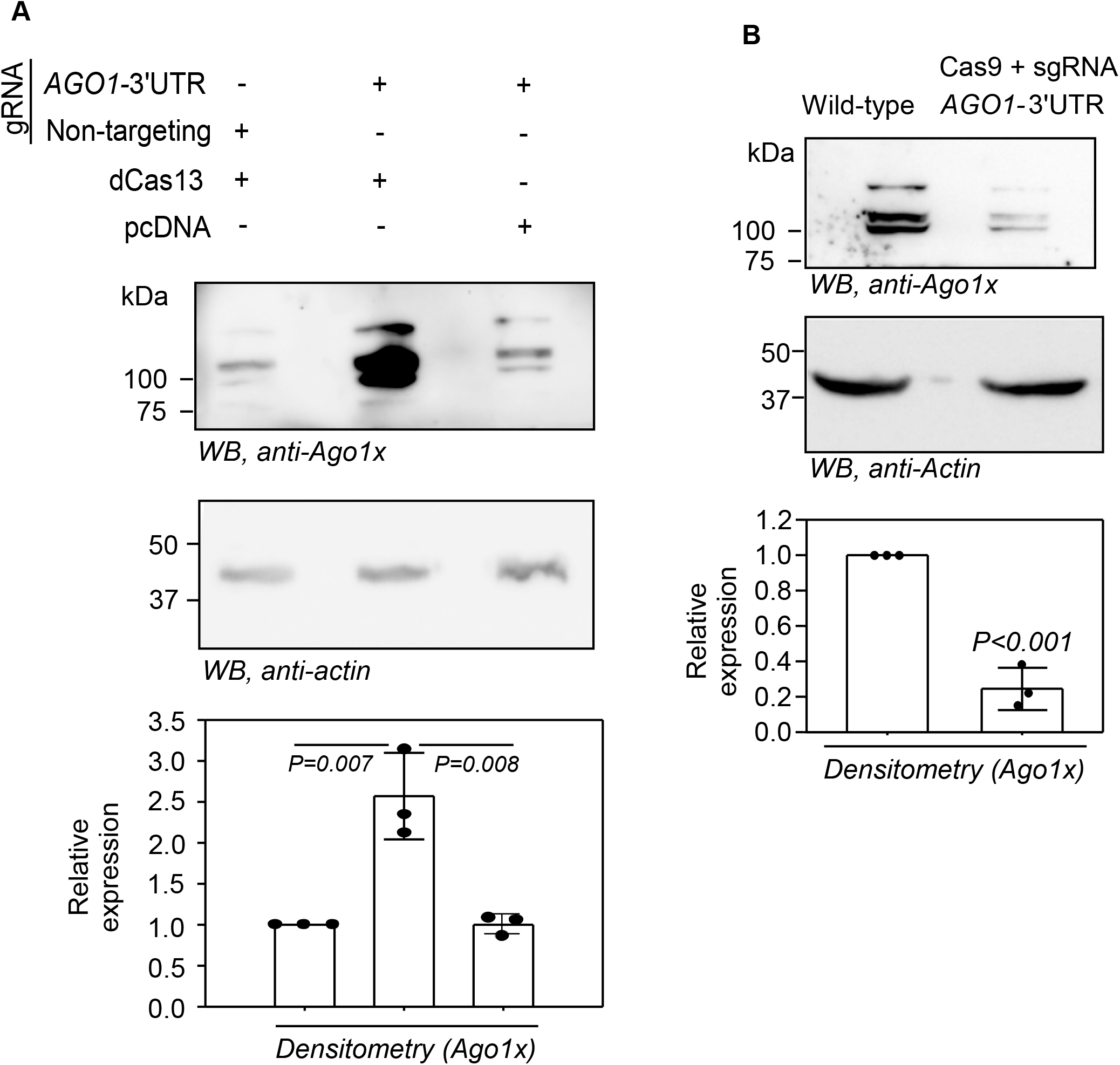
(A) Western blot showing the requirement of both dCas13 and the *AGO1*-3′UTR-targeting gRNA for the increased expression of Ago1x in HEK293 cells. (B) Western blot showing reduced expression of Ago1x in HeLa cells transfected with Cas9 and *AGO1*-3′UTR-targeting sgRNA. Cells were transfected with constructs expressing dCas13 or Cas9 and gRNAs/sgRNAs. Graphs show normalized densitometry values (mean ± sd, N = 3).

